# PFKFB2 Gates a Relationship Between Cardiac Glycolytic Regulation and Electrophysiological Function

**DOI:** 10.64898/2026.01.19.699308

**Authors:** Kylene M. Harold, Harris E. Blankenship, Keaton Minor, Abigail Mulligan, Brooke Loveland, Chi Fung Lee, Michael Kinter, David Kass, Stavros Stavrakis, Michael J. Beckstead, Kenneth M. Humphries

## Abstract

**Background:** The cardiac isoform of phosphofructokinase-2/fructose 2,6-bisphosphatase (PFKFB2) is the heart’s strongest glycolytic regulator but is degraded in the absence of insulin signaling. This makes PFKFB2 loss critical to understand in metabolic heart disease, of which impaired insulin signaling is a hallmark. Prolongation of the QT interval, risk of arrhythmia, and sudden cardiac death are also augmented in metabolic heart disease, raising a question as to whether potential crosstalk between glycolytic dysregulation and electrophysiological dysfunction exists.

**Methods:** We therefore assessed the impact of PFKFB2 loss on cardiac electrophysiology using a cardiomyocyte-specific PFKFB2 knockout mouse model (cKO) and litter-matched controls (CON). To do so, we employed electrocardiography in the fed state and following 12 hours of fasting, examining physiology both at baseline and in the presence of an acute stimulant stress. To further investigate the arrhythmia mechanism, we used patch-clamp electrophysiology and IonOptix Ca^2+^ transient measurements in ventricular cardiomyocytes isolated from CON and cKO hearts.

**Results:** The hearts of cKO mice exhibited prolonged repolarization, marked by QT and action potential duration prolongations. This occurred with impaired Ca^2+^ reuptake and increased spontaneous Ca^2+^ release events in ventricular cardiomyocytes. Ultimately, these changes culminated in ventricular tachyarrhythmia in cKO mice, which was enhanced in the fed relative to the fasted state.

**Conclusion:** These data suggest that in the presence of sufficient glucose availability, cardiac glycolytic dysregulation at the phosphofructokinase nexus is sufficient to promote cardiac electrophysiological instability.

**Clinical Perspective:** *What is Known:*

- Metabolic heart diseases, such as heart failure with preserved ejection fraction and diabetic cardiomyopathy, are associated with heightened risks of arrhythmogenesis and sudden cardiac death.

*What the Study Adds:*

- Here, we show for the first time that PFKFB2 is decreased in human hearts with heart failure with preserved ejection fraction.
- Furthermore, we show that loss of cardiac PFKFB2 is sufficient to promote impaired ventricular repolarization at baseline and ventricular tachyarrhythmia upon stress test.
- This identifies PFKFB2 stabilization and activation as key potential targets in conferring electrophysiological stability in metabolic heart disease.

## Introduction

Metabolic syndromes, heart failure, and arrhythmia have an intricate interplay. The primary presentation of metabolic-associated heart failure, including diabetic cardiomyopathy, is heart failure with preserved ejection fraction (HFpEF), marked by diastolic dysfunction^1^. While HFpEF accounts for approximately half of heart failure cases ^2^ and is increasing in prevalence^3,4^, effective treatment options for HFpEF are limited when compared to heart failure with reduced ejection fraction (HFrEF). This is driven in large part by insufficient understanding of mechanistic underpinnings which could serve as therapeutic targets ^2^.

Among the predominant risk factors for HFpEF are obesity, insulin resistance, poor glycemic control, and diabetes, highlighting the critical relevance of metabolic mechanisms in this disease^5–9^. Of possible contributors to metabolic pathology, a standout candidate is loss of the cardiac isoform of phosphofructokinase-2/fructose 2,6-bisphosphatase (PFKFB2) which regulates the rate-limiting step of glycolysis^10–12^. Our laboratory has previously shown, using both *in vitro* systems and *in vivo* models of diabetes, that PFKFB2 is degraded with the absence or impairment of insulin signaling^12^. Furthermore, we have recently demonstrated that cardiomyocyte-specific knockout of PFKFB2 (cKO) promotes a post-translational modification called O-GlcNAcylation^13^. This modification drives pathology upon chronic upregulation in the heart and is increased in diabetic cardiomyopathy and heart failure of various forms^14–20^. Consistent with this notion, PFKFB2 loss also promotes systolic and diastolic dysfunction, as well as QT prolongation^13^. The latter finding spurred us to further interrogate the cellular changes which underlie this repolarization impairment, as well as the arrhythmogenic potential of a PFKFB2 knockout.

Although atrial arrhythmias are more commonly addressed in metabolically-associated heart disease, the risk of ventricular arrhythmias is also considerably increased in individuals with both HFpEF and diabetic heart disease ^21–23^. Ventricular arrhythmias can lead to sudden cardiac death (SCD), which has an over 8-fold higher incidence in young individuals with diabetes^24^, likely due in part to its association with QT prolongation^25^. Furthermore, SCD is thought to underlie an unexplained overnight mortality, referred to as dead in bed syndrome, which is responsible for 5-6% of mortality in individuals with type 1 diabetes ^26^. This underscores the need for further insight into the mechanisms of ventricular arrhythmia in these metabolic pathologies.

In this study, we show that the cKO model not only displays QT prolongation, but prolongation of the action potential duration, impaired Ca^2+^ reuptake, and increased spontaneous Ca^2+^ release events in individual ventricular myocytes. Furthermore, cKO mice exhibit considerable ventricular tachyarrhythmias which strikingly mimic those observed in a mouse model of severe hyperglycemia under similar stress^27^. This study interrogates a novel and important relationship between glycolytic regulation and electrophysiological dysfunction. This relationship has relevance for various metabolism-related cardiac syndromes and identifies both a pathogenic mechanism and potential therapeutic target for preventing SCD in these more vulnerable populations.

## Methods

### PFKFB2 Measurements and Clinical Correlations

Right ventricular septal endomyocardial tissue was collected from patients with HFpEF and controls without heart failure (unused donor hearts) as previously described ^28,29^ and approved by the institutional review boards at Johns Hopkins University and the University of Pennsylvania. Briefly, biopsies were obtained following informed consent using a standard clinical bioptome (Jawz Bioptome, Argon Medical, Frisco, TX) from patients with HFpEF referred to the Johns Hopkins University HFpEF clinic who met inclusion criteria, or from the University of Pennsylvania biobank (healthy controls).

From patients with HFpEF, clinical information was obtained at the time of first clinic visit. PFKFB2 abundance was measured by proteomic analysis via data independent acquisition-mass spectrometry^28^. Detailed methods and available raw source material are provided in this prior report. Protein abundance values were batch corrected and median-normalized, and resulting fold-change was log_2_ transformed.

### Animals

All animal protocols were approved by the Oklahoma Medical Research Foundation Institutional Animal Care and Use committee. PFKFB2 knockout mice (cKO) and litter-matched control mice (CON) are on a PFKFB2 double floxed background (C57Bl6J). The cKO mice are heterozygous for a cardiomyocyte-specific (myosin-heavy chain 6) Cre, while CON mice are Cre negative. All animals were housed in a 14:10 light:dark cycle (with lights on from 0600 to 2000 hours) and provided a standard ad libitum chow diet. All mice were euthanized by cervical dislocation and cardiac excision under anesthesia (by isoflurane inhalation, either post-electrocardiogram recording or via the drop method).

All ECGs were recorded and mice were sacrificed at approximately 0700 hours, with the exception of fasted mice which were sacrificed at 1900 hours. For the fasted cohort of mice, chow was withheld for 12 hours, from 0700 to 1900 hours (during the light or inactive cycle). ECGs were then acquired at approximately 1900 hours, prior to sacrifice. Metabolic parameters from fasted mice were recently reported^30^.

### In Vivo Electrophysiology

Electrocardiography data were acquired as previously described^13^ and baseline data analyzed as described in supplementary methods. Following five minutes of baseline ECG acquisition, an acute stimulant stress was induced, consisting of intraperitoneal caffeine (100 mg/kg body weight) and epinephrine (1.25 mg/kg body weight) injections. After injection, 20 additional minutes of ECG data were recorded for later arrhythmia analysis. To analyze post-stress ventricular arrhythmia, scores were designated based on most severe rhythm disturbance as previously described^31^, where 1 represents isolated or no premature ventricular complexes (PVCs), 2 represents bigeminy or >10 PVCs per minute, 3 represents a couplet, and 4 represents ventricular tachycardia (VT) which is defined as ≥3 successive PVCs. Bidirectional VT was differentiated from bigeminy when two conditions were met: the net rate during the episode was greater than that in periods of sinus rhythm preceding and following (if resolved) the event, and there was no return to isoelectric baseline between alternating beats.

### Western Blot and Proteomic Analyses in Mouse Hearts

Proteomics and western blots were conducted as described previously ^13,30^, and are also described in detail in the supplemental methods.

### Isolation of Primary Ventricular Myocytes

Mouse hearts were excised, aortas cannulated, and hearts perfused for two minutes with a perfusion buffer containing (in mM): 126 NaCl, 4.4 KCl, 5 MgCl₂·6H₂O, 22 glucose, 20 taurine, 5 creatine, 5 Na pyruvate, 5 NaH₂PO₄, and 10 2,3-butanedione monoxime (BDM) with pH adjusted to 7.4 using NaOH and osmolarity ∼320 mOsm/L. Hearts were then transitioned to perfusion with 47.5 mL of digestion buffer (perfusion buffer supplemented with 2.5 mg/mL collagenase type II and 100 µM CaCl₂). Following digestion, hearts were decannulated into a mixture of 2.5 mL digestion buffer and 5 mL stop buffer (5% calf serum, 100 µM CaCl₂, and 700 µM Na·ATP in perfusion buffer), and mechanically dissociated with tweezers to obtain a single-cell suspension. The cell suspension was then subjected to two sequential washes in stop buffer containing 1.4 mM Na·ATP and increasing CaCl_2_ concentrations (300 µM and 550 µM) which served to gradually raise calcium levels to those in experimental buffers while enriching for viable cells. Finally, cells were moved to a perfusion buffer containing 1 mM CaCl₂ and, in the case of patch clamp experiments, plated for a minimum of 2.5 hours on laminin-coated coverslips before recording.

### Patch Clamp Electrophysiology

Patch pipettes were pulled using standard wall borosilicate glass with tip resistances of 5-10 MΩ. The extracellular solution contained (in mM): 140 NaCl, 5.4 KCl, 1 MgCl₂·6H₂O, 10 HEPES, 5.5 glucose, 1 CaCl₂, and 10 BDM, with pH adjusted to 7.4 using NaOH and osmolarity brought to ∼310 mOsm/L using sucrose. The bath was maintained at 32°C, with extracellular solution perfused at a flow rate of 2 mL/min. The internal pipette solution contained (in mM): 113 K-gluconate, 10 NaCl, 0.5 MgCl₂·6H₂O, 10 HEPES, 5.5 glucose, 0.5 CaCl₂, 10 BDM, 5 K₂·ATP, and 1 EGTA, with pH adjusted to 7.2 using KOH and osmolarity ∼300 mOsm/L.

Current clamp recordings were performed on primary isolated ventricular myocytes in perforated patch using 10 μM β-escin to form large, non-selective pores. This allowed us to achieve a whole cell-like configuration with minimal intracellular dialysis and less mechanical disruption of the membrane. Following approximately 3-5 minutes, access resistance fell below 10-15 MΩ. Cells were then held near −70 mV. To determine rheobase, cells were subjected to a series of 250 ms current steps, increasing in 10 pA increments at 2 Hz. We then provided a 125 ms square current injection corresponding to *rheobase + 50 pA* at 2 Hz from an approximately −70 mV holding potential to elicit an action potential. An average action potential was taken per cell from 100 sweeps. A detailed description of action potential duration quantification at distinct repolarization levels is included in the supplementary methods. Finally, voltage clamp was used to measure capacitance under the same recording conditions.

### IonOptix

After cardiomyocyte isolation, cells were transferred to a bath containing (in mM): 140 NaCl, 5.4 KCl, 1 MgCl_2_·6H_2_O, 10 HEPES, 5.5 glucose, and 1.2 CaCl_2_, pH 7.4. Cells were then protected from light and incubated in 3 µM Fura-2 for 20 minutes. Following incubation, supernatant was removed and cells were allowed to rest in the bath solution without Fura-2 for 15 minutes. Cells were then transferred to the IonOptix chamber where baseline was first recorded for 60 seconds. Cells were next paced at 2.0 Hz and 10.0 V for 100 seconds, then recorded from for 60 more seconds after cessation of pacing to monitor recovery. The IonOptix software was used to obtain metrics of the calcium transients (peak height, time to peak, and time to 50% baseline). Cardiomyocytes were assessed in the last 60 seconds (during the post-pacing recovery period) for spontaneous calcium release events.

### Histology and Masson’s Trichrome Staining

For histological analysis, hearts were excised and immediately placed in 4°C 10% neutral buffered formalin, in which they were kept for 24-48 hours to allow thorough fixation. Hearts were then paraffin embedded vertically, sectioned at 5 µm, and stained with Masson’s trichrome stain. Sections were imaged and quantification performed using Cytation5 (Agilent).

### Statistics

An unpaired student’s *t* test was performed if only 2 groups were present, with the exception of cumulative probability analysis where the Epps-Singleton test was used. When >2 groups were present, a two-way ANOVA was performed with Šidák post hoc test to correct for multiple comparisons. Correlations between PFKFB2 levels and clinical parameters were analyzed via a simple linear regression. Statistical analyses were performed in GraphPad Prism or Python.

### Data Integrity and Availability

Dr. Humphries has access to all study data and takes responsibility for the accuracy of the data analysis and integrity of the data. All raw data and more detailed methods will be made available to other researchers upon request.

## Results

### Patients with HFpEF have decreased cardiac PFKFB2, scalable to heart failure severity and metabolic parameters

PFKFB2 is degraded in the absence of insulin signaling and HFpEF is a predominant presentation of impaired insulin signaling in the heart. We therefore first examined PFKFB2 protein levels in hearts from individuals with HFpEF and healthy controls.

PFKFB2 levels decreased in HFpEF (Figure 1A), and this decrease was scalable to the severity of heart failure as measured by the New York Heart Association (NYHA) classification (Figure 1B). Interestingly, consistent with a close relationship between cardiac PFKFB2 loss and systemic metabolism, PFKFB2 levels were also negatively correlated with body mass index (Figure 1C) and there was a trend toward a negative correlation with hemoglobin A1c (Figure 1D). The latter supports the roles of poor glycemic and insulinemic control as contributing factors to a loss of cardiac PFKFB2 in HFpEF.

**Figure 1.**
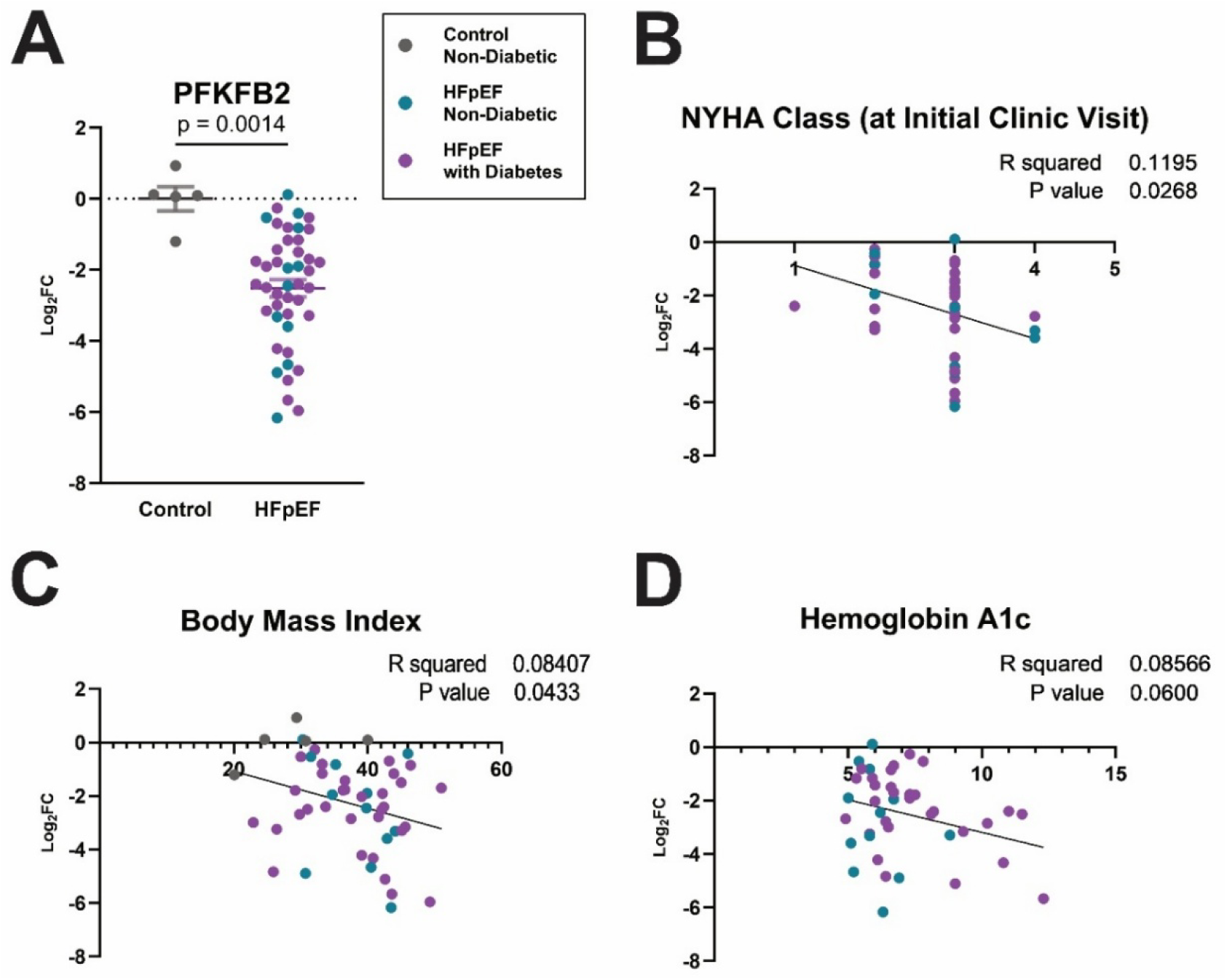
Patients with HFpEF have decreased cardiac PFKFB2, correlated with heart failure severity and systemic metabolism. PFKFB2 was measured by proteomic assay on endomyocardial biopsies from patients with or without Heart Failure with Preserved Ejection Fraction (HFpEF). **A,** PFKFB2 levels are decreased in HFpEF, as measured by Log_2_(fold-change in PFKFB2 levels). **B-D,** Log_2_(fold-change in PFKFB2 levels) is negatively correlated with New York Heart Association Classification (NYHA Class) at initial clinic visit **(B)** and body mass index **(C)**, with a trend toward a negative correlation with Hemoglobin A1C **(D)** by simple linear regression. Grey indicates control (non-failing and non-diabetic; n=5); turquoise indicates HFpEF without diabetes (n=11-12); purple indicates HFpEF with a diabetes diagnosis (n=30-32).

### PFKFB2 cKO promotes ventricular arrhythmogenesis

Given the loss of PFKFB2 in both HFpEF (Figure 1) and diabetes ^12^, we previously generated a cardiomyocyte-specific PFKFB2 knockout mouse model ^13^ and observed that cKO mice died prematurely at a mean age of 9 months with no prior signs of decline, suggesting an arrhythmic etiology. While echocardiographic changes did not explain this mortality, QT prolongation and a sudden nature of the mortality spurred us to further interrogate potential arrhythmogenic propensity. To do so, we treated mice for 20 minutes with an acute pharmacologic stress of caffeine and epinephrine, sufficient to induce arrhythmia only in the presence of electrophysiological instability. Doing so led to a large number of premature complexes (PVCs) in cKO mice, but very few in CON mice (on average, 79.6 in cKO as compared to 2.2 in CON hearts over a 20-minute period, p=0.0027; Figure 2A, B). Additionally, the cKO hearts spent approximately 23% of the stress period in ventricular tachyarrhythmia, whereas the CON mice did not exhibit these more severe ventricular arrhythmias (Figure 2A, C). Finally, we utilized a ventricular arrhythmia score to denote the most severe ventricular rhythm disturbance to occur in each mouse ^31^. Almost all CON mice exhibited either no arrhythmia or only had isolated PVCs, with just two mice exhibiting couplets. Conversely, all but two cKO mice exhibited ventricular tachycardia (VT), with the two that did not still exhibiting some degree of high PVC burden or couplets (Figure 2D). This culminated in an increased ventricular arrhythmia score in cKO mice (p <0.0001; Figure 2D). This is of particular interest given the association of VT with HFpEF and diabetes, diseases marked by cardiac PFKFB2 loss.

**Figure 2.**
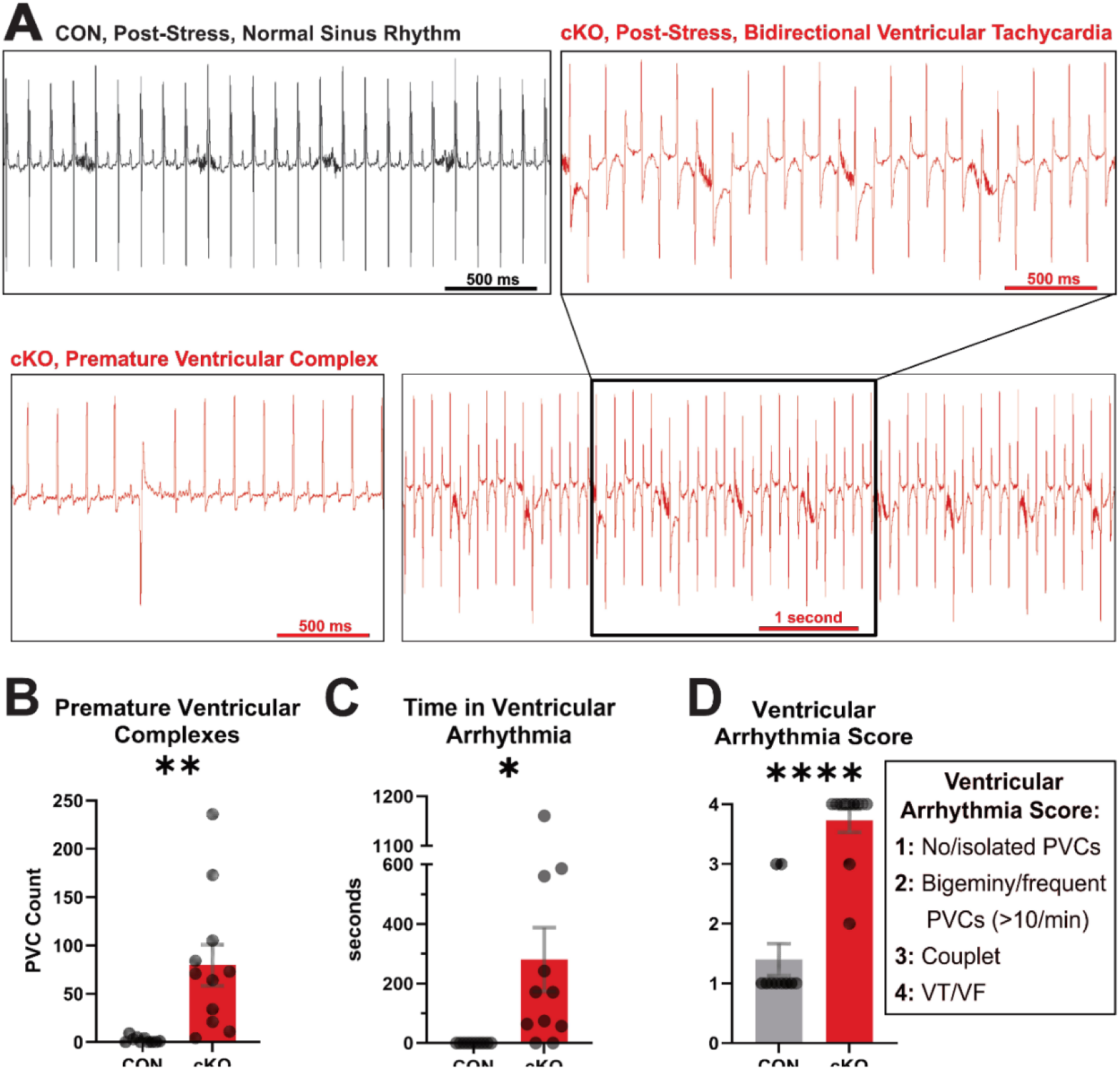
Loss of cardiac PFKFB2 promotes ventricular arrhythmogenesis. Mice were stressed with caffeine and epinephrine. **A,** Representative traces post-stress showing a control in normal sinus rhythm (top left), a PFKFB2 knockout exhibiting a premature ventricular complex (bottom left), and a PFKFB2 knockout exhibiting bidirectional ventricular tachycardia (right). **B-D,** There is an increase in premature ventricular complexes **(B)**, time in ventricular tachyarrhythmia **(C)**, and ventricular arrhythmia score **(D)** in PFKFB2 knockout hearts (n=11) relative to controls (n=10) over a 20 minute stress period. cKO indicates PFKFB2 cardiomyocyte-specific knockout and CON indicates litter-matched controls. \**P*≤0.05, \*\**P*≤0.01, \*\*\*\**P*≤0.0001 by unpaired student’s *t* test.

### Loss of cardiac PFKFB2 promotes delayed repolarization at the whole heart and single cell levels

We next began investigating possible substrates for this arrhythmia. We first examined structural substrate in the form of fibrosis via trichrome staining. As expected, with bidirectional ventricular tachycardia being a triggered arrhythmia, there was no increase in fibrosis in cKO ventricles (Supplemental Figure 1). However, consistent with our previous report showing that cKO hearts demonstrate an eccentric hypertrophy phenotype by echocardiography^13^, capacitance was greater in isolated cardiomyocytes from cKO relative to CON ventricles, as determined using patch clamp electrophysiology (Supplemental Figure 2).

Another possible substrate for arrhythmia that is commonly associated with VT is delayed repolarization, as manifested by QT interval prolongation. Beyond signifying increased propensity for fatal ventricular arrhythmias, prevalence of QT prolongation is considerably increased in diabetes ^27^. This prolongation is also a well-accepted predictor of mortality and SCD propensity in individuals with diabetes ^25,32,33^ and HFpEF^34^, as well as in the general population^35,36^. Consistent with our previous study^13^, we observed QT prolongation in cKO hearts relative to CON (p=0.0032; Figure 3A-B), which was sustained when corrected for RR interval (Figure 3C-D). This appears largely attributable to impairment of early repolarization, here marked by an effectively abolished J wave amplitude (p=0.0007; Figure 3E), which is compensated for temporally later in repolarization (p= 0.0074; Figure 3F). Remarkably, the profound reduction in J wave amplitude and QT prolongation remarkably phenocopy models of diabetic hyperglycemia^27^ despite the cKO mice in this study being normoglycemic.

**Figure 3.**
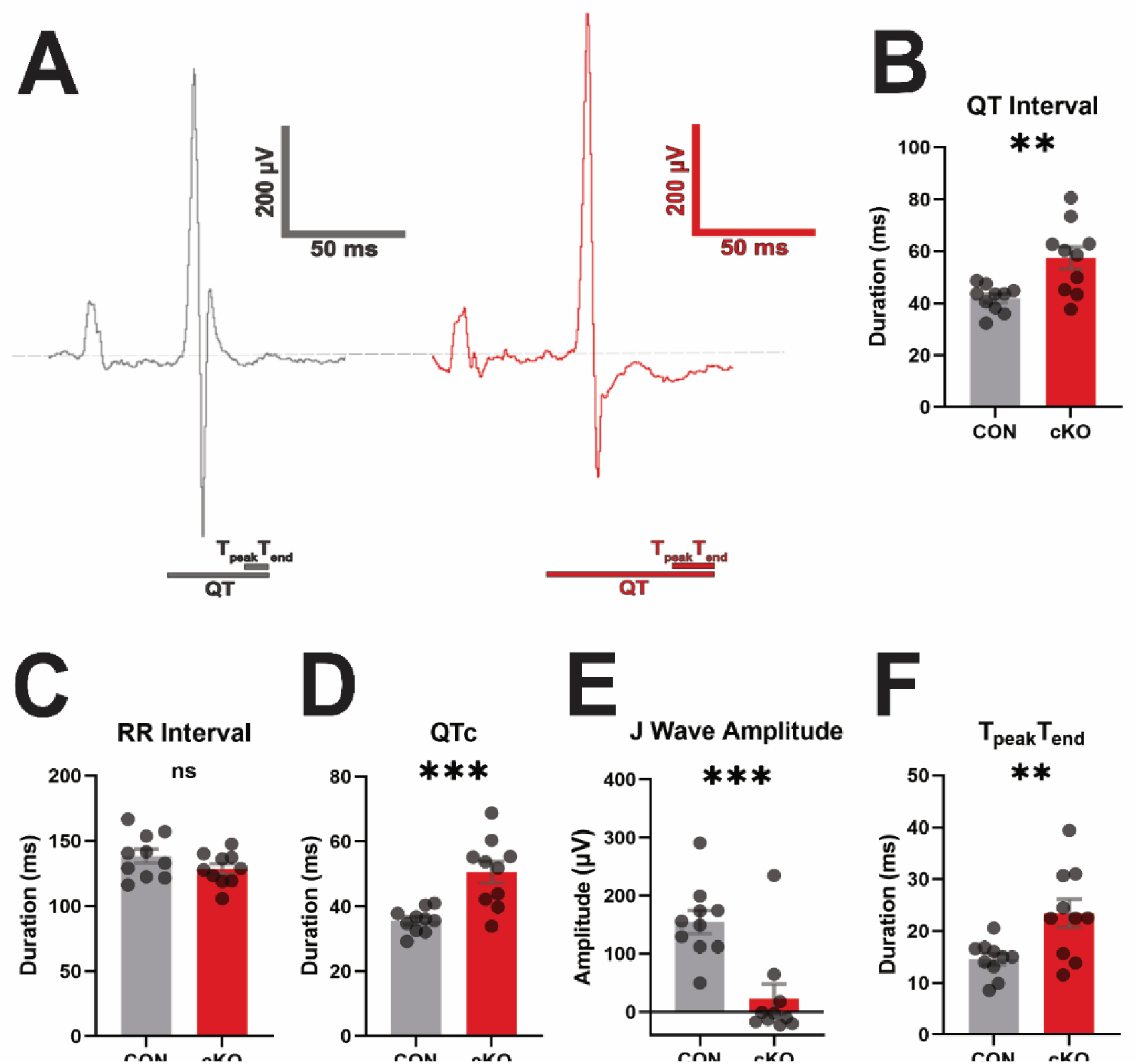
PFKFB2 cKO mice exhibit changes in ventricular repolarization. Electrocardiography was assessed at baseline by surface ECG**. A,** Example traces from control (grey) and PFKFB2 knockout (red) hearts are shown. **B,** QT interval is prolonged in PFKFB2 knockout hearts. **C-D,** RR interval is unchanged by genotype **(C)** so corrected QT interval (QTc) is also prolonged **(D)**. **E-F,** J wave amplitude is decreased **(E)** and the interval from the peak to the end of the T wave (T_peak_T_end_) is prolonged in knockout hearts **(F)**. cKO indicates PFKFB2 cardiomyocyte-specific knockout and CON indicates litter-matched controls. n=10 per group. ns = not significant, \*\**P*≤0.01, \*\*\**P*≤0.001. by unpaired student’s *t* test.

We next sought to further assess this delayed repolarization in cKO hearts through patch clamp electrophysiology experiments using primary ventricular cardiomyocytes isolated from adult CON and cKO hearts. Analysis of experiments performed in current clamp showed prolongation of action potential duration (APD) in cKO cardiomyocytes at multiple phases of repolarization (Figure 4A-C). This was demonstrated by prolongation at both 10% (APD 10) and 90% (APD 90) of repolarization (Figure 4B, C). In agreement with the loss of J wave amplitude (Figure 3E), prolongation of APD 10 (Figure 4B) suggests an impairment of the earliest phase of repolarization. This is typically attributable to either changes in the transient outward repolarizing K+ current (I_to_) or changes in gating kinetics of the depolarizing voltage-gated Na^+^ current (I_Na_). Consistent with the latter, when we assessed action potential kinetics (Figure 4D), we found that both the maximal instantaneous rates of depolarization (Figure 4E) and repolarization (Figure 4F) were impaired, implicating changes in I_Na_.

**Figure 4.**
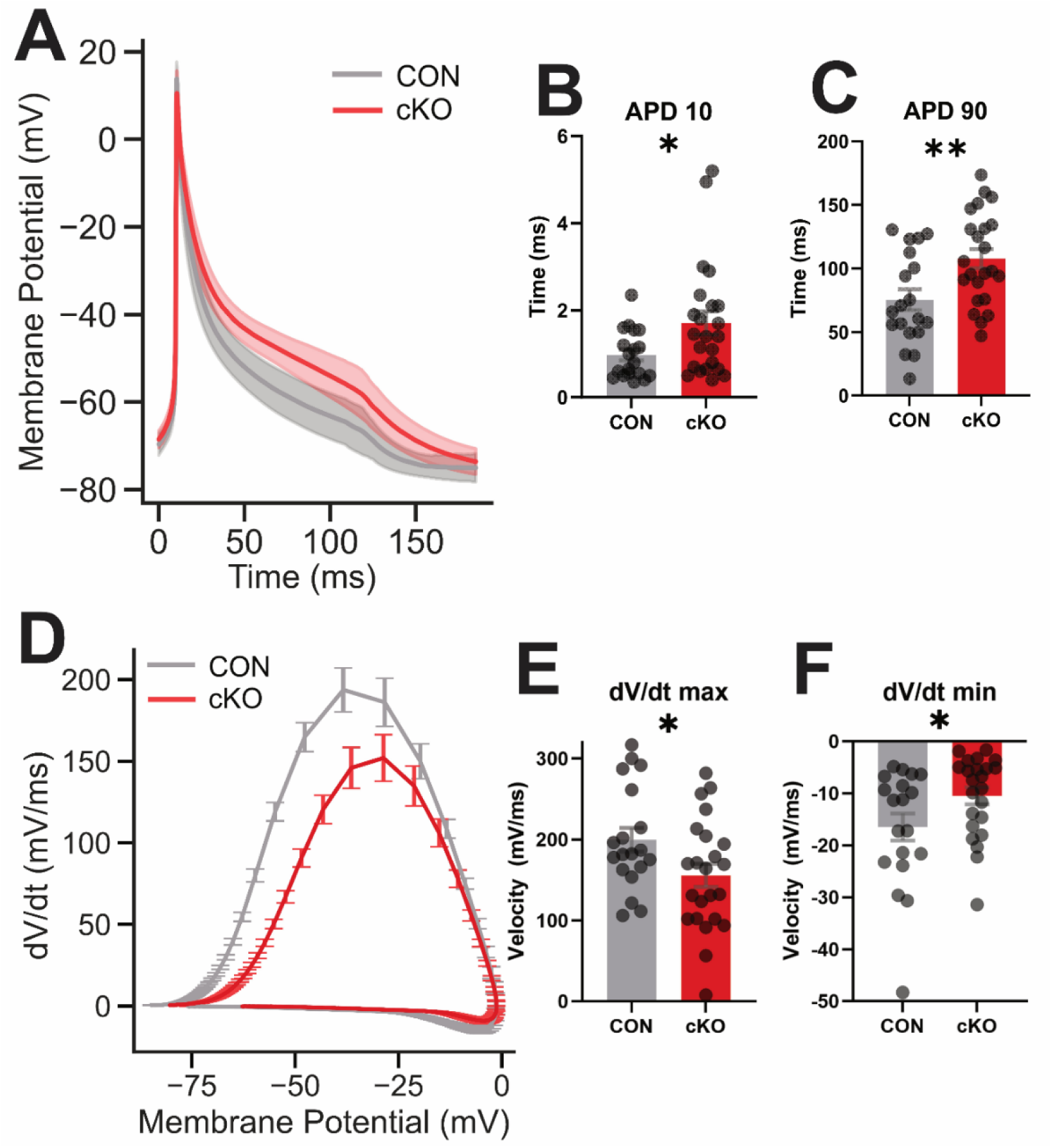
Cardiomyocytes from cKO ventricles exhibit distinct action potential kinetics, marked by impaired repolarization. Action potentials were recorded from isolated primary ventricular cardiomyocytes in perforated patch and current clamp conditions. **A,** Averaged action potential waveforms across all cells (dark red, grey) with SEM (lighter red, grey shading). **B-C,** Action potential duration at 10% **(B)** and 90% **(C)** of repolarization is prolonged in PFKFB2 knockout cardiomyocytes. **D,** Averaged phase plane plot across all cells. **E-F,** maximum and minimum rate of change in membrane potential is impaired in PFKFB2 knockout cardiomyocytes (n=23 cells from 3 mice) relative to controls (n=19 cells from 5 mice). cKO indicates PFKFB2 cardiomyocyte-specific knockout and CON indicates litter-matched controls. \**P*≤0.05, \*\**P*≤0.01 by unpaired student’s *t* test.

### Ca^2+^ handling is impaired in PFKFB2 cKO hearts

Changes in intracellular Na^+^ can alter Ca^2+^ handling via the Na^+^/Ca^2+^ exchanger (NCX), potentially promoting changes in repolarization. Furthermore, changes in Ca^2+^ handling are among the most implicated mechanisms underlying bidirectional VT. Therefore, we next utilized the IonOptix system to assess intracellular Ca^2+^ transient properties in isolated primary cardiomyocytes from CON and cKO mice. After 60 seconds of baseline measurement, we field-paced ventricular cardiomyocytes for 100 seconds at 2.0 Hz and 10.0 V, then monitored their recovery for spontaneous Ca^2+^ release events for 60 seconds (Figure 5A). Upon analysis of transients acquired during the pacing period, we found that there was no difference in peak height or time to peak of the Ca^2+^ transients (Figure 5B-D; Supplemental Figure 3A-B). Though, the time to 50% baseline was prolonged in cKO relative to CON cardiomyocytes (Figure 5B,E; Supplemental Figure 3C), suggesting an impairment in calcium reuptake or clearance from the cardiomyocyte. Reuptake of cytosolic Ca^2+^ into the sarcoplasmic reticulum is mediated in the cardiomyocyte by the sarco/endoplasmic reticulum Ca^2+^ ATPase type 2 (SERCA2), which is inhibited by phospholamban (PLB)^37^. Phosphorylation of PLB at Ser16 or Thr17 in response to protein kinase A (PKA) or Ca^2+^/calmodulin-dependent protein kinase II (CAMKII), respectively, releases its inhibition of SERCA2^38,39^. While there were no significant differences in abundance of SERCA2 (Figure 5F) or PLB (Figure 5G), we did observe a surprising increase in phosphorylated PLB (Ser16/Thr17; Figure 5H). However, due to a numeric increase in total PLB, the ratio of p-PLB to PLB was not different (Figure 5I). This leaves ambiguity surrounding whether PLB or an alternate mechanism was impairing Ca^2+^ clearance from the cytosol.

**Figure 5.**
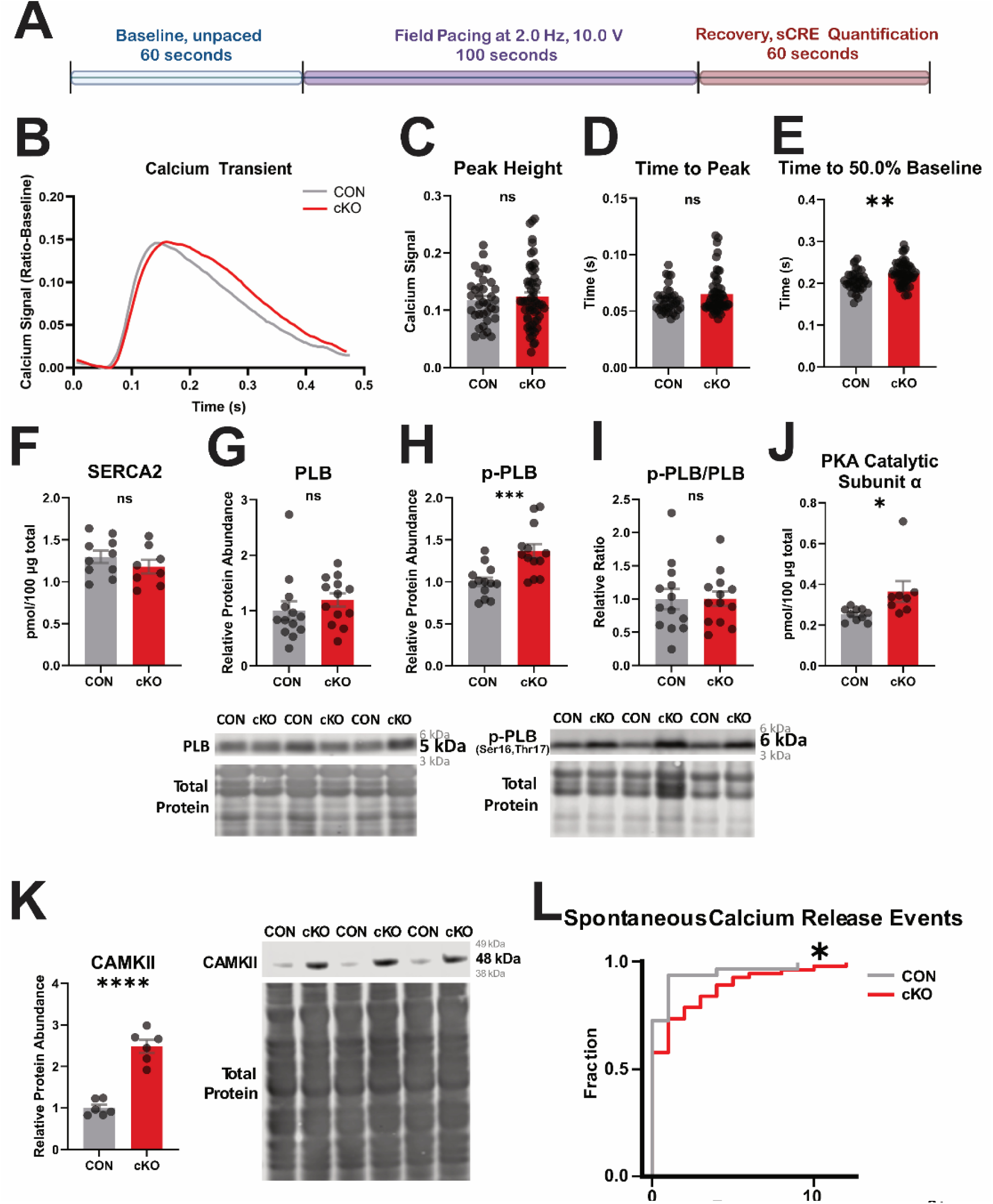
Absence of PFKFB2 in ventricular myocytes promotes impaired Ca^2+^ reuptake and spontaneous Ca^2+^ release events. **A,** Schematic of IonOptix protocol used to acquire data shown in panels B-E and L from primary ventricular cardiomyocytes. **B,** Example traces generated using a rolling ball average of 3-4 cells and subtraction of the lowest value. **C-E,** Peak height, time to peak, and time to 50% baseline are quantified from isolated ventricular PFKFB2 knockout and control cardiomyocytes (n=35-57 cells from 4-5 mice). **F,** Sarcoplasmic/endoplasmic reticulum ATPase 2 (SERCA2) is measured by proteomics (n=8-10 per group). **G-H,** Phospholamban (PLB; **G)** and phosphorylated phospholamban (p-PLB (Ser16/Thr17); **H)** are measured by western blot (n=13 per group). **I,** the ratio of p-PLB/PLB from **G-H** is taken (n=13 per group). **J,** Protein Kinase A (PKA) catalytic subunit alpha is measured by proteomic analysis(n=8-10 per group). **K,** Ca^2+^/Calmodulin-activated protein kinase II (CAMKII) was measured by western blot (n=6 per group). **L,** Spontaneous Ca^2+^ release events, acquired from primary ventricular myocytes in the final phase of the IonOptix protocol, displayed as cumulative probability (n=33-57 cells from 4-5 mice). cKO indicates PFKFB2 cardiomyocyte-specific knockout and CON indicates litter-matched controls. ns = not significant, \**P*≤0.05, \*\**P*≤0.01, \*\*\**P*≤0.001, \*\*\*\**P*≤0.0001 by unpaired student’s *t* test **(A-K)** or Epps-Singleton test **(L)**.

In investigating the mechanism of PLB phosphorylation, we found only a modest increase in PKA abundance (Figure 5J) with no change in PKA substrate phosphorylation (Supplemental Figure 4). However, the abundance of CAMKII, which is activated in response to elevated cytosolic Ca^2+^, was increased 2.48-fold in cKO mice relative to controls (p<0.0001; Figure 5K). One implication of CAMKII hyperactivity is an increase in spontaneous Ca^2+^ release events via Ryanodine Receptor (RyR) sensitization ^17,40^. Indeed, when we measured spontaneous Ca^2+^ release events from the final phase of the pacing protocol, we found that by cumulative probability analysis, cKO cardiomyocytes exhibited a greater number of events than CON (p=0.0153; Figure 5L). This demonstrates the presence of a potential trigger for the bidirectional VT or other ventricular rhythm disturbances.

### Electrophysiological implications of PFKFB2 loss are partially fed state-dependent

Finally, we have previously shown that cKO hearts exhibit an increase in the post-translational modification O-linked N-acetylglucosamine (O-GlcNAc)^13^. This modification is supported by activity of the hexosamine biosynthesis pathway, an ancillary pathway which branches from glycolysis immediately upstream of the step regulated by PFKFB2. We have recently proposed that the phosphofructokinase nexus acts as a valve, such that when PFKFB2 is degraded and canonical glycolysis inhibited, metabolites are instead shunted to the hexosamine biosynthesis pathway, driving O-GlcNAcylation^30^.

We have also recently shown that this is fed-state dependent, with O-GlcNAcylation decreasing in the fasted state, likely due to a decrease in glucose substrate availability for the hexosamine biosynthesis pathway^30^. Because chronic O-GlcNAcylation promotes pathology in the heart, as well as Ca^2+^ mishandling and SCD predisposition^14,17,41,42^, we hypothesized that the decrease in O-GlcNAcylation secondary to fasting would confer some degree of electrophysiological protection in the cKO heart. To test this, we fasted mice for 12 hours prior to the ECG recording with acute caffeine and epinephrine stress (Figure 6A) and compared these mice to those stressed in the same manner in the fed state (data carried from Figure 2). We found that there was no difference in ventricular arrhythmia score with fasting, suggesting no difference in incidence of most severe arrhythmia (Figure 6B). However, considerably more PVCs were observed in the fasted cKO relative to fed cKO mice (Figure 6C), and conversely, considerably less time was spent in ventricular arrhythmia in the fasted relative to fed cKO mice (Figure 6D). This suggests that while the fasted mice experienced the same ventricular tachyarrhythmias, they more quickly returned to normal sinus rhythm. Similarly, the higher PVC burden in fasted mice reflects that many of these events occurred in isolation or couplets and did not progress to sustained VT, whereas in fed mice, PVCs more often converted to VT or prolonged ventricular arrhythmia episodes due to reduced electrophysiological stability.

**Figure 6.**
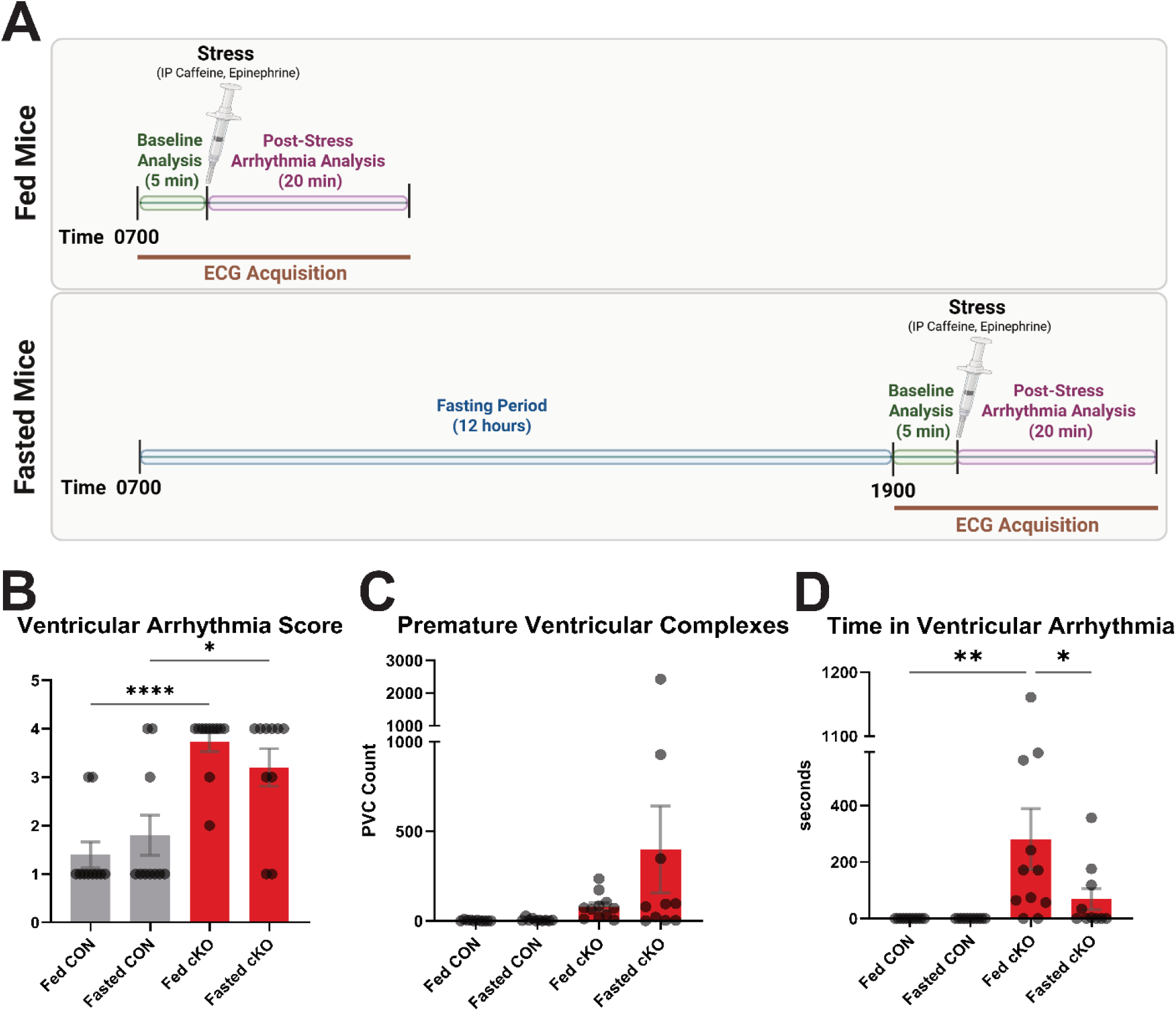
Electrophysiological instability in cKO hearts is partially fed state dependent. **A,** Schematic feeding, stress, and acquisition time points in the fed and fasted states. **B,** Ventricular Arrhythmia score, as defined in figure 2D. **C-D,** Abundance of premature ventricular complexes **(C)** and time in ventricular arrhythmia **(D)** over a 20 minute period. Note: fed CON and cKO data for reference was drawn from Figure 2. Analysis was performed by 2-way ANOVA with Šidák test for multiple comparisons, testing and correcting for comparisons which differed by only one variable. n=10-11 per group. If not denotated, comparison is not significant. *P≤0.05, **P≤0.01, ****P≤0.0001.

## Discussion

Prior investigation into the relationship between glycolysis and ion handling in the heart has predominantly focused on how ion pumps and channels utilize or are sensitive to glycolytically-derived ATP. The heart primarily relies on fatty acid oxidation for ATP production. However, a functional compartmentalization has been demonstrated, such that ATP-producing glycolytic enzymes are tightly co-localized with sarcoplasmic reticulum or plasma membrane pumps and channels. This promotes their preferential response to, or use of, locally-produced glycolytic ATP. For example, the Na^+^/K^+^ pump and SERCA respond more robustly to blockade of glycolysis or supplementation of glycolytic precursors than to inhibition of oxidative phosphorylation or supplementation of exogenous ATP^43,44^. Similarly, following exposure to a system that consumed ATP, ATP-sensitive K^+^ channels responded to glycolytic intermediates but not mitochondrial metabolic substrates^45,46^. Finally, when glycolysis is inhibited and pyruvate is used in place of glucose, functionally replacing glycolysis with glucose oxidation, alternans are observed in cardiac myocytes. A plausible explanation involves the RyR, which is responsible for Ca^2+^ release from the sarcoplasmic reticulum. It has been suggested that decreased ATP production by glycolytic enzymes could impair the localized activity of kinases such as PKA that phosphorylate RyR^47^. This would subsequently impair RyR activation and recovery, making RyR less available on alternating beats, and promoting alternans. Collectively, these studies highlight the importance of local glycolytic ATP production in maintaining ionic homeostasis and orchestrating coordinated ionic movement across essential membranes.

However, our previous metabolic investigations in the cKO model demonstrate increased abundance of GLUT1, the heart’s insulin-independent glucose transporter, as well as increased glycolytic enzyme abundance and downstream glucose oxidation^13^. These data suggest that the capacity for upregulated glycolytic ATP production exists, implying that inhibition of glycolytic ATP production itself does not drive the electrophysiological differences observed in the cKO model. Instead, these data point toward glycolytic regulation, rather than glycolytic activity or ATP production, as a key mechanism of electrophysiological dysfunction in the cKO model.

Another factor to which the relationship between glucose metabolism and electrophysiology has been attributed is altered glycemic status. For example, while streptozotocin (STZ)-treated animal models demonstrate hyperglycemia, the intracellular downstream effects of hypoinsulinemia, such as PFKFB2 loss, are often overlooked. Notably, the cKO mice exhibit many changes which parallel or even phenocopy these hypo-insulinemic and subsequently hyperglycemic models^27^, yet remain normoglycemic in the fed state^13,30^. This indicates that intracellular glucose handling, rather than extracellular glucose, drives dysregulation in the cKO heart.

After excluding two of the most intuitive links between glucose and ion mishandling in the cKO heart, we considered additional aspects of intracellular glucose utilization that could promote electrophysiological remodeling and arrhythmogenesis. The post-translational modification O-GlcNAcylation emerged as a standout candidate because the hexosamine biosynthesis pathway, which provides substrate for O-GlcNAcylation, branches from glycolysis immediately upstream of the phosphofructokinase regulatory nexus. O-GlcNAcylation is increased in cKO mice, suggesting that when flux through canonical glycolysis is blunted, upstream substrate overflows into the hexosamine biosynthesis pathway. Consistent with this paradigm, the increase in O-GlcNAcylation observed in cKO mice is ameliorated by 12 hours of fasting^30^. Because the heart takes up very little glucose in the fasted state, this finding suggests that decreased substrate for the HBP can alone reduce O-GlcNAcylation, minimizing the impact of PFKFB2 loss. In line with the idea that activation of ancillary pathways in the fed state links phosphofructokinase nexus impairment to electrophysiological dysfunction, we show here that 12 hours of fasting also significantly decreased ventricular tachyarrhythmia duration (Figure 6D).

O-GlcNAcylation is highly implicated in electrical stability and Ca^2+^ handling alterations. The relationship between O-GlcNAcylation and Ca^2+^ regulation is especially nuanced due to its bidirectionality^48,49^. Studies conducted in glucose-deprivation have shown a Ca^2+^ dependence of O-GlcNAcylation^49^. O-GlcNAcylation in turn impacts intracellular Ca^2+^ handling via multiple context-specific mechanisms. For example, increased O-GlcNAcylation has been shown to attenuate Ca^2+^ overload in ischemically-stressed cardiomyocytes^50^. Yet, conversely, studies conducted under normoxic conditions demonstrated that O-GlcNAc modification can also impair Ca^2+^ reuptake via downregulation of SERCA2^51–53^ or O-GlcNAcylation of PLB^54^. The latter is thought to promote PLB-mediated inhibition of SERCA2 by blocking PLB phosphorylation^54^. However, we observed unchanged SERCA protein levels and increased PLB phosphorylation in the cKO model.

PLB phosphorylation is primarily mediated by PKA or the Ca^2+^ effector CAMKII. In a seminal study investigating the relationship between O-GlcNAcylation and Ca²⁺ handling, the Bers laboratory demonstrated that O-GlcNAcylation modifies and activates CaMKII in hyperglycemic conditions^17^. This modification of CAMKII promotes Ca^2+^ mishandling, including increased spontaneous Ca^2+^ release events, and ultimately arrhythmogenesis^17,41^. Further, hyperglycemia-mediated CAMKII O-GlcNAcylation broadly downregulates K^+^ channel expression, impairing repolarization reserve and ultimately promoting electrophysiological instability^55^. It is therefore plausible that a mechanism of dysfunction in cKO hearts is O-GlcNAcylation of CaMKII that compounds its significantly increased abundance in the cKO heart (Figure 5K). RyR is a key target of CAMKII and is also modified by O-GlcNAcylation. Phosphorylation by CAMKII^56,57^ or O-GlcNAcylation^58^ both sensitize RyR, promoting sarcoplasmic reticulum Ca^2+^ release to the cytosol. In excess, this can lead to Ca^2+^ overload and spontaneous Ca^2+^ release events. Furthermore, O-GlcNAc can impact less commonly considered Ca^2+^ handling mechanisms in the heart such as store operated Ca^2+^ entry via modification of STIM1^59^. Taken together, O-GlcNAcylation impacts Ca^2+^ handling through numerous mechanisms.

Finally, another shared substrate of both O-GlcNAcylation^60^ and CAMKII^61,62^ is Nav1.5. Interestingly, modification by either of these factors promotes the late Na^+^ current (I_Na,L_)^60,61,63,64^. I_Na,L_ involves a shift in Nav1.5 inactivation kinetics, such that a sub-population of Nav1.5 channels fails to inactivate before, or reopens during, the plateau phase of the action potential. The resulting sustained inward Na^+^ current can indirectly elevate cytosolic Ca²⁺ through reverse-mode NCX^65–67^. I_Na,L_, secondary to O-GlcNAcylation or CAMKII-mediated phosphorylation, has been shown to serve as a QT-prolonging and ventricular arrhythmogenic mechanism in models of diabetes^60^ and HFpEF^68^. Thus, impaired phosphofructokinase nexus regulation may drive Ca²⁺-dyshomeostasis through O-GlcNAcylation and CaMKII hyperactivity, via an array of convergent and distinct pathways.

These O-GlcNAcylation and CAMKII-mediated mechanisms have direct implications for metabolic heart disease beyond pre-clinical models. For example, the aforementioned increase in O-GlcNAcylation of CAMKII has been established both in a rat model of diabetic hyperglycemia and in human patients with type 2 diabetes (T2D)^17^. Clinically, these molecular changes parallel the elevated arrhythmic risk observed in diabetes. Matched controls have an 18% lower risk for ventricular tachycardia or fibrillation than patients with T2D by hazard ratio^22^. Moreover, the risk of sudden cardiac arrest is heightened even further in type 1 diabetes than T2D, underscoring that severe metabolic dysregulation markedly amplifies arrhythmogenic vulnerability. Beyond the described Ca^2+^ dysregulation that likely supports arrhythmic trigger, repolarization abnormality also contributes to the substrate. QTc prolongation is observed in up to 44% of patients with diabetic heart disease, considerably increasing the risk for SCD^25,69^.

Heart failure and diabetes have a stark interplay^8^, culminating in their comorbidity. The Reykjavik study showed a ∼12% prevalence of heart failure in individuals with diabetes as compared to ∼3% in individuals with normal glucose regulation^70^. It is therefore anticipated that electrophysiological abnormalities observed in diabetes would also manifest in HFpEF. Indeed, there is increased prevalence of ventricular tachycardia in HFpEF patients^21,23^, and those which experience VT incidents have significantly longer QTc^21^. Even among individuals with hypertrophic cardiomyopathy, HFpEF is associated with higher VT incidence relative to those without heart failure, highlighting a role for non-structural mechanisms^71^. While VT in HFpEF is often non-sustained, the presence of non-sustained VT has been shown to be associated with 3.4-fold higher odds of mortality over a median three year follow up^72^. Drug-induced arrhythmias further highlight this vulnerability. Observational work has shown that QT-prolonging drugs such as dofetilide or sotalol have augmented effects on QTc prolongation^73^ and VT risk^74^ in patients with HFpEF relative to those without HF. Sudden death remains a primary mode of death in HFpEF, with reduced conduction velocity, delayed repolarization, and ion-channel remodeling contributing to arrhythmic risk in this population. Thus, QTc prolongation and ventricular arrhythmias are clinically important features of HFpEF and warrant further investigation.

Metabolic-associated heart failure and ventricular arrhythmia share common comorbidities such as obesity^75^, which is of interest as BMI is negatively correlated with PFKFB2 abundance (Figure 1C). These conditions also share various underlying mechanisms, making their comorbidity unsurprising^76^. Such mechanisms include increased cytosolic Ca^2+^ load, metabolic insufficiency, O-GlcNAcylation, and other metabolic processes. Each of these underpinnings promotes both impaired relaxation and membrane electrical instability. However, shared upstream processes which could be driving them remain elusive. Here, we identify PFKFB2 loss as a single molecular change that could contribute to many of these shared mechanisms, affecting both mechanical and electrical function in the heart and promoting arrhythmogenesis.

### Limitations

The use of mouse models of electrophysiology is inherently limited in that repolarization rate, activation patterns, and path length differ considerably between human and murine hearts. Therefore, a specific arrhythmogenic presentation, such as bidirectional ventricular tachycardia, may not be directly reproduced across species under comparable pathological conditions. However, underlying mechanisms of arrhythmia, such as the substrate provided by QT prolongation, or changes in Ca^2+^ handling, are generally conserved across models of disease. While data must be interpreted with these limitations in mind, arrhythmogenesis in mouse models nonetheless signifies considerable electrical instability.

### Future directions

Much of this work could be expanded to consider atrial electrophysiological implications, especially given the relative predominance of atrial arrhythmias in metabolic heart disease. Additionally, further mechanistic studies should investigate the impacts of direct O-GlcNAcylation inhibition on electrical (patho)physiology in the cKO heart.

Alternatively, these mechanistic studies could target the pentose phosphate or polyol pathways which also utilize metabolites upstream of the phosphofructokinase nexus in glycolysis. Finally, future studies should address the protective impact of PFK-2 activity on cardiac electrophysiology in models of diabetes and HFpEF, initially using the Glyco^Hi^ kinase-dominant PFK-2 overexpression model, and eventually moving to pharmacologic approaches. A PFKFB2 isoform-specific activator could hold great value in both mitigating the impacts of ancillary pathway activation and restoring metabolic flexibility, while minimizing pathologic off-target effects in other tissues.

## Acknowledgements

We thank the Oklahoma Medical Research Foundation Imaging Core for their assistance with preparation of trichrome-stained heart sections. We also thank Vivek P. Jani, Aleksandra Binek, and Virginia S. Hahn for their roles in generating and analyzing the human proteomics data.

## Supplemental Materials

Supplemental Methods

Supplemental Figures 1-4

## Abbreviations

APD: Action Potential Duration
BSA: Bovine Serum Albumin
CAMKII: Ca^2+^/Calmodulin-Dependent Protein Kinase II
cKO: PFKFB2 Cardiomyocyte-Specific Knockout
CON: Control
HFpEF: Heart Failure with Preserved Ejection Fraction
HFrEF: Heart Failure with Reduced Ejection Fraction
NCX: Na^+^/Ca^2+^ Exchanger
NYHA: New York Heart Association
O-GlcNAc/O-GlcNAcylation: O-linked N-Acetylglucosamin(e/ation)
PFKFB2: Phosphofructokinase-2/Fructose 2,6-Bisphosphatase 2
PKA: Protein Kinase A
PLB: Phospholamban
PVCs: Premature Ventricular Complexes
QTc: Corrected QT Interval
RyR: Ryanodine Receptor
SCD: Sudden Cardiac Death
SDS: Sodium Dodecyl Sulfate
SERCA2: Sarcoplasmic/Endoplasmic Reticulum Ca^2+^ ATPase 2
STZ: Streptozotocin
TBS: Tris Buffered Saline
T2D: Type 2 Diabetes
VT: Ventricular Tachycardia

---

**Table 1.**
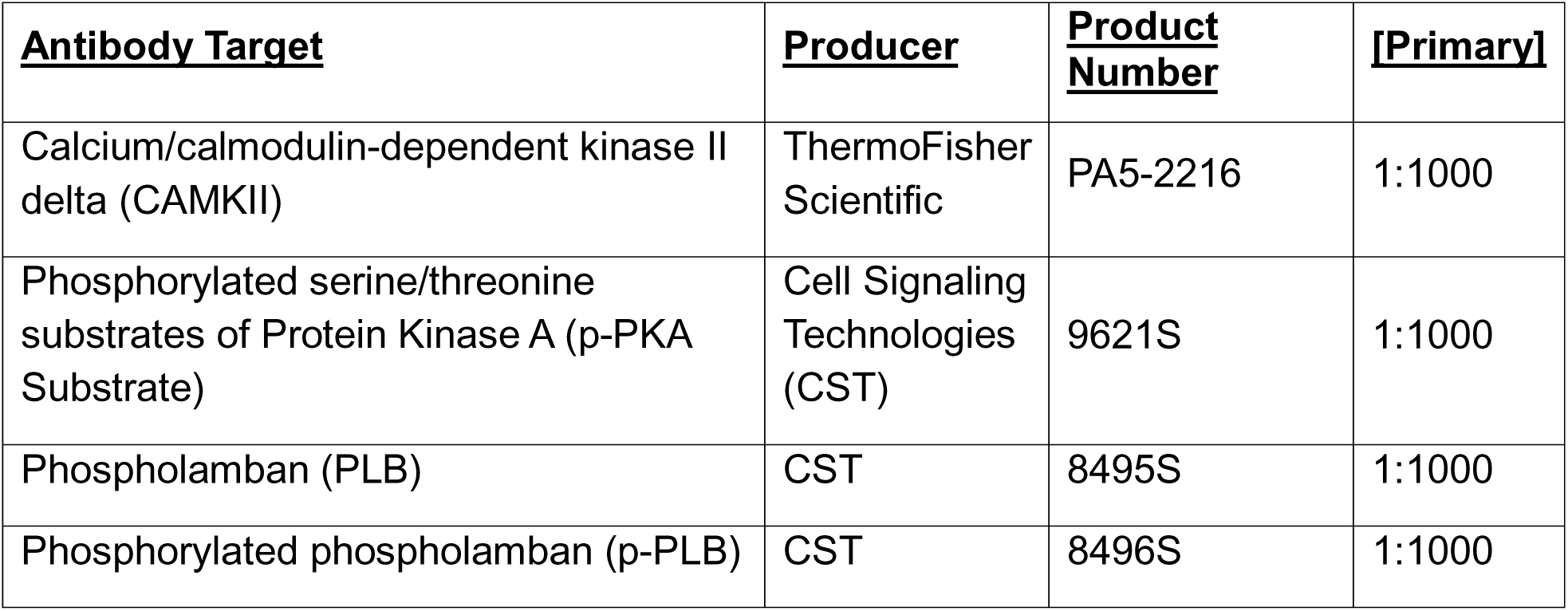
Antibodies Used in Western Blot Experiments.

| **<u>Antibody Target</u>** | **<u>Producer</u>** | **<u>Product Number</u>** | **<u>[Primary]</u>** |
| --- | --- | --- | --- |
| Calcium/calmodulin-dependent kinase II delta (CAMKII) | ThermoFisher Scientific | PA5-2216 | 1:1000 |
| Phosphorylated serine/threonine substrates of Protein Kinase A (p-PKA Substrate) | Cell Signaling Technologies (CST) | 9621S | 1:1000 |
| Phospholamban (PLB) | CST | 8495S | 1:1000 |
| Phosphorylated phospholamban (p-PLB) | CST | 8496S | 1:1000 |

