## Supplemental Material for "PFKFB2 Gates a Relationship Between Cardiac Glycolytic Regulation and Electrophysiological Function"

**Western Blot and Proteomic Analyses in Mouse Hearts**

Hearts were homogenized on ice in 210 mM mannitol, 70 mM sucrose, 5 mM 3-(N-morpholino)propanesulfonic acid, 1 mM EDTA, with pH of 7.4, after which they were centrifuged at 4°C and 550×g for 5 minutes to produce a working homogenate.

For proteomic analysis, a mixture of 100 µL working homogenate, 100 µL 1% sodium dodecyl sulfate (SDS), and 20 µL 10% SDS was heated. Based on protein quantification, 100 µg protein was used for analysis. An additional 100 µL 1% SDS was added, along with 1 µg bovine serum albumin (BSA) as an exogenous internal standard, and the samples were heated at 70 °C before overnight precipitation with acetone. The precipitate was reconstituted to 1 µg/µL in Laemmli sample buffer and run on an SDS-PAGE gel, after which each lane (sample) was cut into smaller pieces, washed, reduced, alkylated, and digested with 1 µg trypsin overnight at room temperature. Peptides were extracted from the gel in 50% acetonitrile. Extracts were dried by Speedvac and reconstituted in 200 µL 1% acetic acid.

For selected reaction monitoring, previously validated assay panels were used to measure respective groups of proteins, with most assays monitoring 2 peptides per protein. Fifteen data points across 30 second chromatographic peaks were acquired using a cycle time of 2 seconds. Liquid chromatography involved linear gradient elution from 2%B to 50%B in 60 minutes.

For data independent acquisition, a 20 m/z window worked from m/z 350 to 950. The orbitrap was operated at 17,500 and a full scan spectrum was acquired at 70,000 each cycle. These conditions give 7 to 8 data points across 30 second chromatographic peaks. A large group of internally developed assays, each using 2 to 3 peptides that have been previously validated, was used for analysis.

Skyline was used to locate and integrate proper chromatographic peaks for both methods. Retention times are predicted based on calibration using BSA and trypsin peptides and calculations determine the protein response based on a geomean of the two monitored peptides. All data are normalized to internal standard (BSA).

For western blot, working homogenate was heated at 95°C with 25% lithium dodecyl sulfate, 20 mM dithiothreitol, and 1% Halt protease and phosphatase inhibitor. In the case of western blots for membrane-bound proteins, lysis buffer was also added to samples prior to heating. For most proteins, samples were run at 200 V for 45 minutes and transferred using a semi-dry transfer system at 30 V for 60 minutes. In the case of large proteins (>100 kDa), samples were run at 200 V for 75 minutes and transferred using the semi-dry system at 30V for 120 minutes, with the chamber placed on ice throughout. Membranes were stained for total protein and imaged using the LI-COR system (Odyssey CLx). They were then probed with their respective primary antibodies (Table 1) by rocking overnight at 4°C and then for an additional hour at room temperature. Following 3 washes (10 min each at room temperature) in Tris-Buffered Saline (TBS) with 1% tween, blots were incubated in secondary at 1:3000 (IRDye 800CW, LI-COR) with rocking for 60 minutes at room temperature. Washes were then repeated with the final wash consisting of TBS without tween. Blots were imaged again using the LI-COR system and analyzed along with previous total protein stain images to which they were normalized (Image Studio, Version 5.2).

**In Vivo Electrophysiology Acquisition and Baseline Electrocardiography Analysis**

To measure in vivo cardiac electrophysiology (lead II), we placed needle electrodes subcutaneously in the right front and both hind limbs. Mice were anesthetized with 2.0% isoflurane for induction and 1.0-1.5% for maintenance. Data were acquired at a sampling rate of 4 kHz using an AD instruments data acquisition system (PowerLab) for digitization, equipped with a Bio Amplifier. Filter settings included a high pass filter at 0.3 Hz and a low pass filter at 1 kHz. For baseline ECG analysis, beat averaging view was employed, averaging increments of 20 beats, including only sinus beats, aligned at the peak of the R wave. The cumulative average of parameters measured from these averaged beats over a total time of 2 minutes was taken. The corrected QT interval (QTc) was calculated using a modified Bazett's formula for mice[31], QTc = QT/(sqrt(RR/100)).

**Action Potential Duration Calculation at Distinct Repolarization Levels From Current Clamp Data**

APD values at 10% and 90% repolarization (APD 10 and APD 90) were calculated using the following method. For each action potential, the peak voltage ($V_{peak}$) and baseline voltage ($V_{baseline}$) were first determined. The baseline voltage was defined as the membrane potential at the first point where dV/dt exceeded 0.5 mV/ms during the upstroke phase. Repolarization voltage thresholds were then calculated as $V_{APD_x} = V_{baseline} + (1 - x/100) \times (V_{peak} - V_{baseline})$ where $x$ represents the percent repolarization (10 or 90). APD was defined as the time interval between the peak of the action potential and the first point during repolarization where the membrane potential fell to $V_{\mathrm{APD}_x}$. If the voltage threshold was not reached during repolarization, APD was measured to the point of minimum voltage during the repolarization phase.

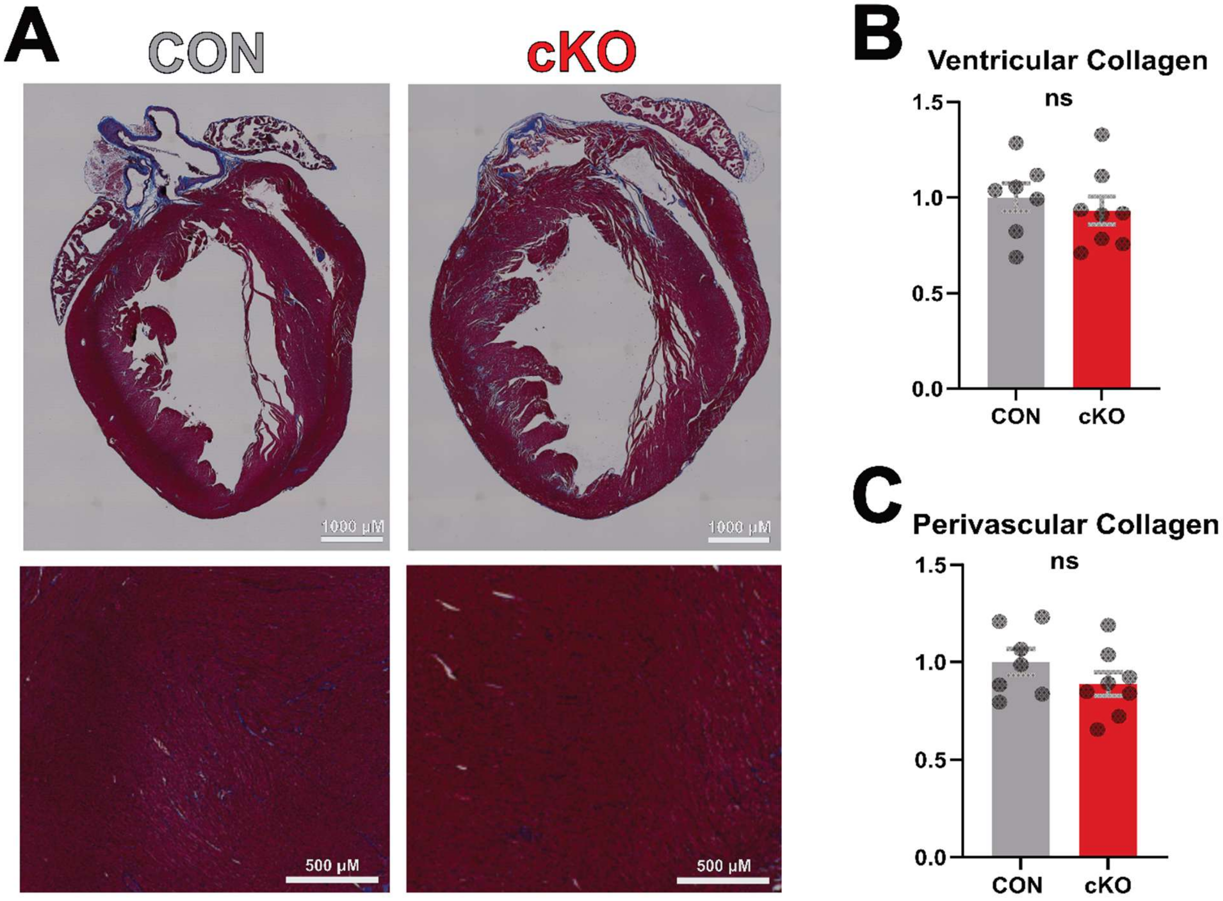


**Supplement 1. There is no difference in fibrosis by Masson's trichrome staining in cKO relative to control hearts.**

**A,** Example images of trichrome stained vertical heart sections. **B-C,** Relative quantification of fibrosis in the whole ventricular wall **(B)** or perivascular area **(C)**. cKO indicates PFKFB2 cardiomyocyte-specific knockout and CON indicates litter-matched controls. n=7-8 per group. ns = not significant by unpaired student's *t* test.

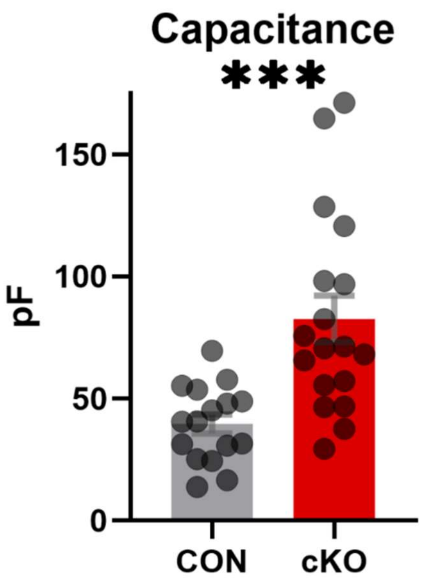


**Supplement 2. Capacitance is larger in cKO than CON cardiomyocytes.**

Capacitance was measured in primary cardiomyocytes isolated from cKO and CON ventricles. pF indicates picofarads. cKO indicates PFKFB2 cardiomyocyte-specific knockout and CON indicates litter-matched controls. n=16-18 per group. ****P*≤0.001 by unpaired student's *t* test.

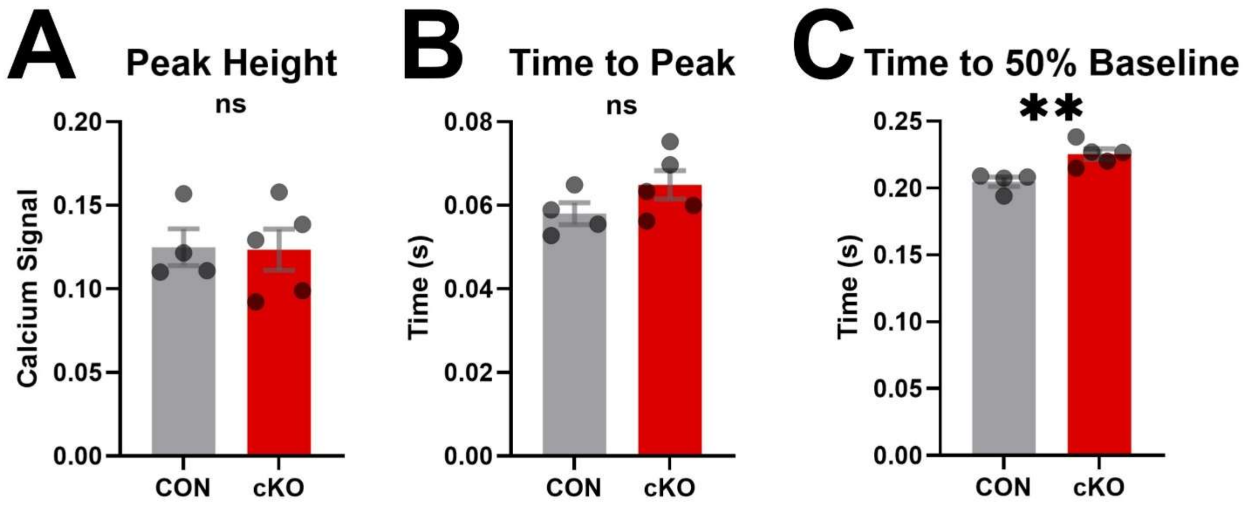


**Supplement 3. Animal-level averages of $Ca^{2+}$ data corroborate individual cell averages.**

Data shown in figure 5 are averaged by animal from which ventricular cardiomyocytes were isolated, with each animal shown as a single data point. CON indicates control (n=4) and cKO indicates knockout (n=5). ns indicates not significant (p>0.05), ***P*≤0.01.

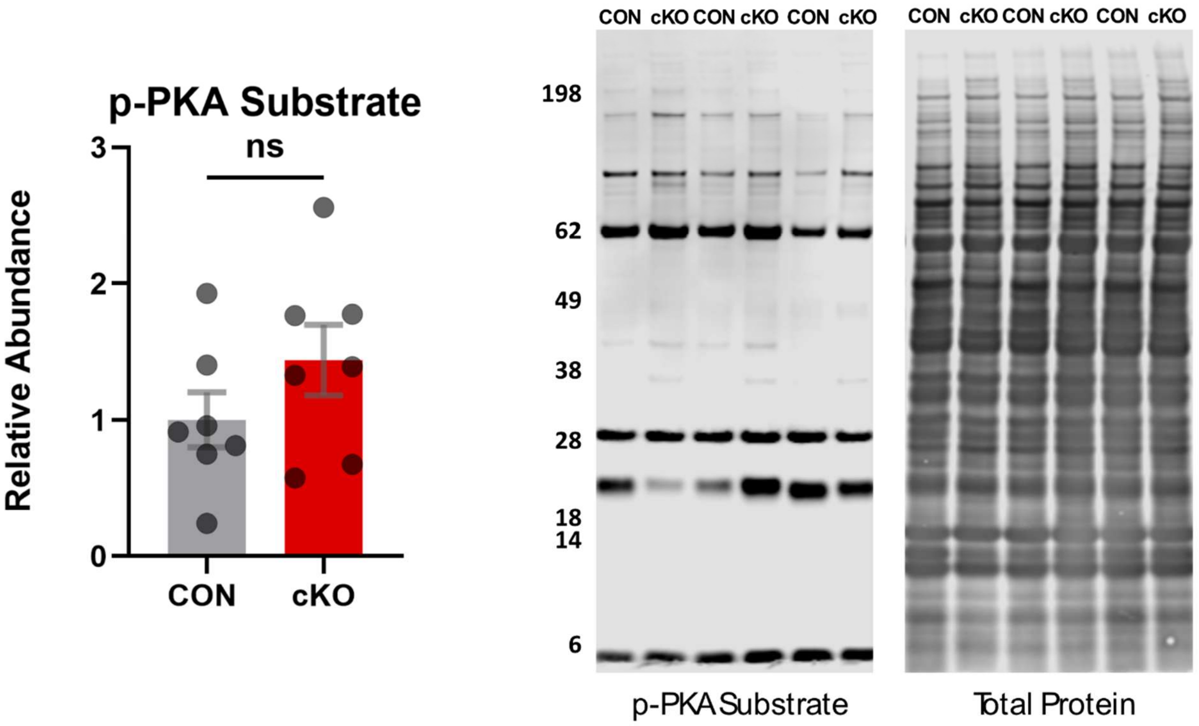


**Supplement 4. There is no significant difference in abundance of phosphorylated PKA substrate in cKO relative to CON hearts.**

A pan-specific antibody targeting phosphorylated PKA substrates was utilized for a western blot of working homogenate from the ventricles. cKO indicates PFKFB2 cardiomyocyte-specific knockout and CON indicates litter-matched controls. n=7 per group. ns = not significant by unpaired student’s *t* test.
